# Structural analysis of genetic variants of the human tumor suppressor Palb2 coiled-coil domain

**DOI:** 10.1101/2024.03.19.585655

**Authors:** Pothula Purushotham Reddy, Apurva Phale, Ranabir Das

## Abstract

Homologous recombination (HR) is an obligate pathway to repair DNA lesions and maintain genome integrity. The tumor suppressor Palb2 is an integral part of the HR pathway that functionally connects BRCA proteins at the site of DNA damage. The N-terminal coiled-coil domain in Palb2 switches from an inactive to an active complex during HR. In the inactive form, Palb2 forms homodimers, and during HR, it assumes the active state, forming a heterodimeric complex with BRCA1. However, the structural details of the human Palb2 coiled-coil domain are unknown. About 600 missense variants have been reported in Palb2, and several are in the coiled-coil domain. The structure-function relationship of such variants is poorly understood, which challenges genetic counseling. Here, we report the solution structure of the human Palb2 coiled-coil domain, which forms an antiparallel homodimer. We then use the structure to study the effect of a few well-characterized missense mutations on the fold and interactions of the Palb2 coiled-coil domain. The structural impact of mutations correlates excellently with their efficiency in homologous recombination, signifying that the approach can be exploited to study other genetic variations in the Palb2.

## Introduction

Double-strand DNA breaks (DSB) are severe lesions that introduce discontinuity in chromatin. The two major DSB repair pathways are non-homologous end joining (NHEJ) and homologous recombination (HR). NHEJ is an error-prone mechanism by which DNA ends are joined, whereas HR is a high-fidelity mechanism that uses the sister chromatid as a homologous DNA template for repair. To repair a DNA break, HR starts by creating single-stranded 3’ overhangs through DNA end resection. These overhangs are then coated with the ssDNA-binding protein RPA (Replication protein A) to safeguard them from degradation and creating non-specific secondary structures. The BRCA2-RAD51 protein complex and BRCA1 are recruited to the RPA-coated ssDNA. The BRCA2 protein assists in creating a nucleoprotein filament by loading RAD51 onto the ssDNA, which enables efficient repair of the DNA break. Mutations in genes encoding homologous recombination proteins in mammals, such as BRCA1 and BRCA2, which are associated with breast and ovarian cancer susceptibility, have been linked to developmental abnormalities and tumorigenesis. However, some families affected by these conditions have no pathogenic mutations in either BRCA1 or BRCA2 genes. The discovery of *Palb2* (Partner and Localizer of BRCA2) has revealed a crucial gene associated with susceptibility to breast cancer, Fanconi Anemia, and other cancers^7^.

The *Palb2* gene encodes an 1186 amino acid long protein consisting of several domains, including the N-terminal coiled-coil domain, ChaM (chromatin association motif) domain, and C-terminal WD40 domain. Palb2 is an essential bridging molecule between BRCA1 and BRCA2 and an integral component of the BRCA complex. The ChaM domain located in the center of the protein is responsible for localizing Palb2 to the chromatin. The C-terminal ring-like B-propeller structure of the WD40 domain of Palb2 interacts with BRCA2^1^. The WD40 domain also interacts with other essential proteins for HR like the RAD51, RAD51C, and RNF168^2,3^Human Palb2 exists either as a homodimer with minimal activity (OFF state) or as a heterodimer with BRCA1 (ON state)^4^. The N-terminal coiled-coil domain homodimerizes, and the same domain interacts with BRCA1 to form a heterodimer during HR^5^. Overexpression of the Palb2 coiled-coil domain reduces RAD51 foci formation during HR, indicating a competition between Palb2 homo- and heterodimers and suggesting that the molecular switch from Palb2 homodimer to heterodimer can regulate HR^4^. In the OFF state, the coiled-coil domain forms an oligomer^6^. In the ON state, a heterodimer is formed between the BRCA1 coiled-coil domain and the Palb2 N-terminal coiled-coil domain^5^. The mouse Palb2 coiled-coil domain suggest that it form antiparallel dimer^22^. However, the structure of the human Palb2 coiled-coil domain is unknown, and the atomistic interactions that regulate the switch from the ON to OFF state are poorly understood.

Similar to BRCA1 and BRCA2, Palb2 is a tumor suppressor. Biallelic mutation in Palb2 can cause Fanconi anemia, and monoallelic mutations can increase susceptibility to breast cancer, ovarian cancer, and pancreatic cancer^7^. Multiple Variants of Uncertain Significance (VUS) are present in Palb2^8^. Most VUS exist within the N-terminal coiled-coil or the C-terminal WD-40 domain (Figure 1A). HeLa cells expressing various VUS were tested for their PARPi (olaparib) sensitivity. L35P, K18R, R37H, and Y28C showed varying degrees of sensitivity. L35P showed the highest sensitivity, with a survival rate below 20%, suggesting severely impaired HR^9^. Functional analysis of a few missense mutations in the Palb2 coiled-coil domain showed varying effects on HR^8–10^. L35P impairs HR by more than 90%. Y28C, R37H, and L24S mutations impair HR by >60% (Figure 1B). It is unclear how these mutations affect the Palb2 homodimer and its switch to heterodimer. The ability to effectively analyze missense mutations in the Palb2 coiled-coil domain and predict the resulting impact on Palb2 function is crucial in evaluating other variants of unknown significance, which plays an integral role in genetic counseling.

**Figure 1.**
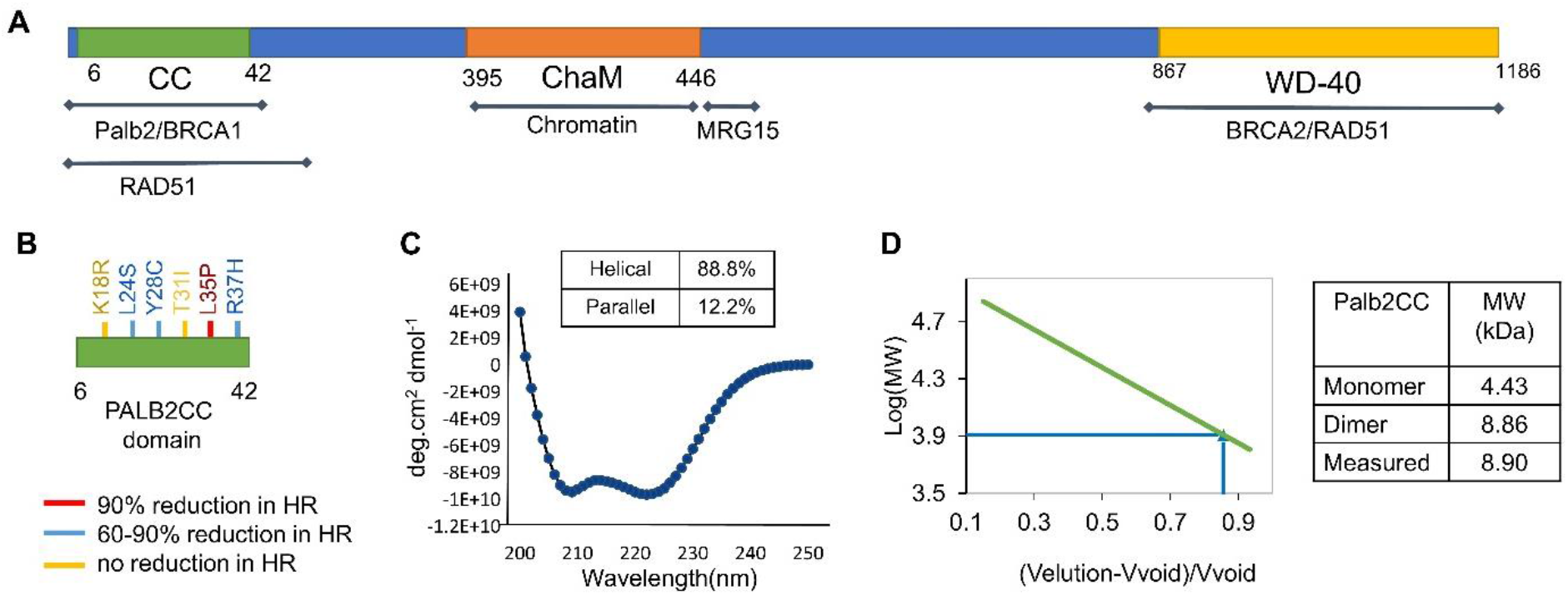
(A) Schematic representation of Palb2 protein and its structural domains along with the known binding sites of its interactors. The coiled-coil domain spans amino acids 6-42. (B) A few of the identified variants of the Palb2 in the coiled-coil domain were followed up in this study. (C) The CD spectrum of the Palb2cc domain. The helical and parallel beta-sheet propensities obtained from fitting the CD data are mentioned at the top. (D) The size of the Palb2cc domain is determined by analytical size exclusion chromatography. The plot of molecular weight standards against the elution volume is on the left. Expected monomer, dimer molecular weights, and the measured molecular weight are given on the right.

Here, we study the N-terminal coiled-coil domain of human Palb2 (Palb2cc). Biophysical studies confirmed that Palb2cc has a helical secondary structure and forms a homodimer. We then solve the solution structure for the Palb2cc, which reveals an antiparallel homodimer. The dimer interface consists of multiple hydrophobic contacts spanning the length of the helix and an aromatic-aromatic interaction at the center. Molecular dynamics simulations suggest that each monomer is unstable due to the exposed hydrophobic residues, and the dimer structure buries them to stabilize the protein. We then investigated the impact of specific genetic variants on Palb2cc homodimer and Palb2cc/BRCA1cc heterodimer structures. Our detailed structural analysis demonstrates how these variants modulate the stability of Palb2cc and how this destabilization correlates with Palb2’s HR efficiency. This study provides a framework for analyzing other VUS within the Palb2cc domain and predicting their impact on Palb2’s activity during DNA repair.

## Experimental procedures

### Protein Expression and Purification

A chimeric construct of Palb2 coiled-coil domain (residues 5-41) with N-terminal His-tagged GB1 was cloned in the pET3a vector. For CNBr cleavage, a methionine residue was introduced at the N-terminus of the Palb2 coiled-coil domain. This construct was overexpressed in BL21-DE3 cells in LB media for unlabeled protein and ^13^C, ^15^N enriched M9 media for isotope-labeled protein. The cells were grown till OD_600_ reached 0.6 and were induced with 1mM IPTG for 4 hours. The cells were harvested, re-suspended in 50 mM Na_2_HPO_4_, 300 mM NaCl, and 20 mM imidazole (pH 8), and lysed by sonication. The lysate was centrifuged, and the supernatant was incubated with pre-equilibrated Ni^2+^-NTA beads. The beads were washed, and the protein was eluted with 20%, 40%, and 60% elution buffer (50 mM Na_2_HPO_4_, 300 mM NaCl, 500 mM Imidazole). Further purification was carried out by a gel filtration column (HiLoad 16/600 Superdex75 pg, GE). The pure fractions were pooled, concentrated to 6 mg/ml, and cleaved by CNBr to remove the N-terminal tags. The cleaved protein was separated with reverse phase column chromatography (RPC) using a gradient of up to 60% methanol with 0.1% TFA. The pure fractions containing Palb2cc collected from RPC were concentrated and lyophilized.

### Circular Dichroism (CD)

Palb2cc, L35P-Palb2cc, and S10E, S29E-Palb2cc peptides were synthesized from Lifetein tenchologies. The peptides were dissolved in 50 mM Na_2_HPO_4_, 150 mM NaCl pH 6.5, and degassed before measurement. CD measurements were performed on a Jasco CD spectrophotometer in a 1 mm path-length quartz cuvette. Each wavelength scan was performed with 1 nm increments and bandwidth from 250 to 200 nm. The thermal melt experiment was collected with a 1°C increment every 2 minutes. All CD data were averaged for ten seconds at each measurement, and the buffer signal was subtracted. Reversibility (often between 70 and 96%) was determined by re-measuring the signal at 222 nm and 20°C.

### Analytical Size Exclusion Chromatography (ASEC)

The proteins were dissolved in 50 mM Na_2_HPO_4_, 150 mM NaCl, 5 mM DTT, and pH 6.5. ASEC (HiLoad 30/300 Superdex75 pg, GE) column was used to calculate the molecular mass of each protein.

### NMR Data Collection and Structure Calculation

NMR data for resonance assignments and structure determination were collected at 25°C using Bruker AVANCE III 600 and 800 MHz NMR spectrometers equipped with 5 mm cryoprobes. Complete ^1^H, ^13^C, and ^15^N resonance assignments for Palb2cc were determined using conventional triple-resonance (HNCO, CBCACONH, HNCACB, HCCCONH3D3, HCCCONH3D2, and HCCHTOCSY) NMR methods. Standard 3D ^15^N-NOESY-HSQC and ^13^C-NOESY-HSQC experiments with a mixing time of 100ms were used to get NOE restraints. NMR data Processing and Analysis were done using NMRpipe^11^ and NMRfam_Sparky^12^, respectively. The dimer interface on Palb2cc was identified using ^13^C filtered NOESY experiments on a 1:1 sample of ^13^C, ^15^N enriched, and unlabeled (natural abundance) protein. The Palb2cc homodimer structure was calculated by using CYANA version 2.1^13^ and refined in XPLOR-NIH. The dihedral angle constraints obtained from TALOS+4 were used for structure calculations. In addition, the intrachain and interchain NOE data were included. The backbone dihedral angles and bond lengths of the final converged structures were evaluated by the Molprobity^14^ and PSVS^15^ suite of programs. The NMR constraints and refinement statistics are provided in Table 1. The structure and data is deposited with PDB id 8YAP and BMRB id 36647.

**Table 1.**
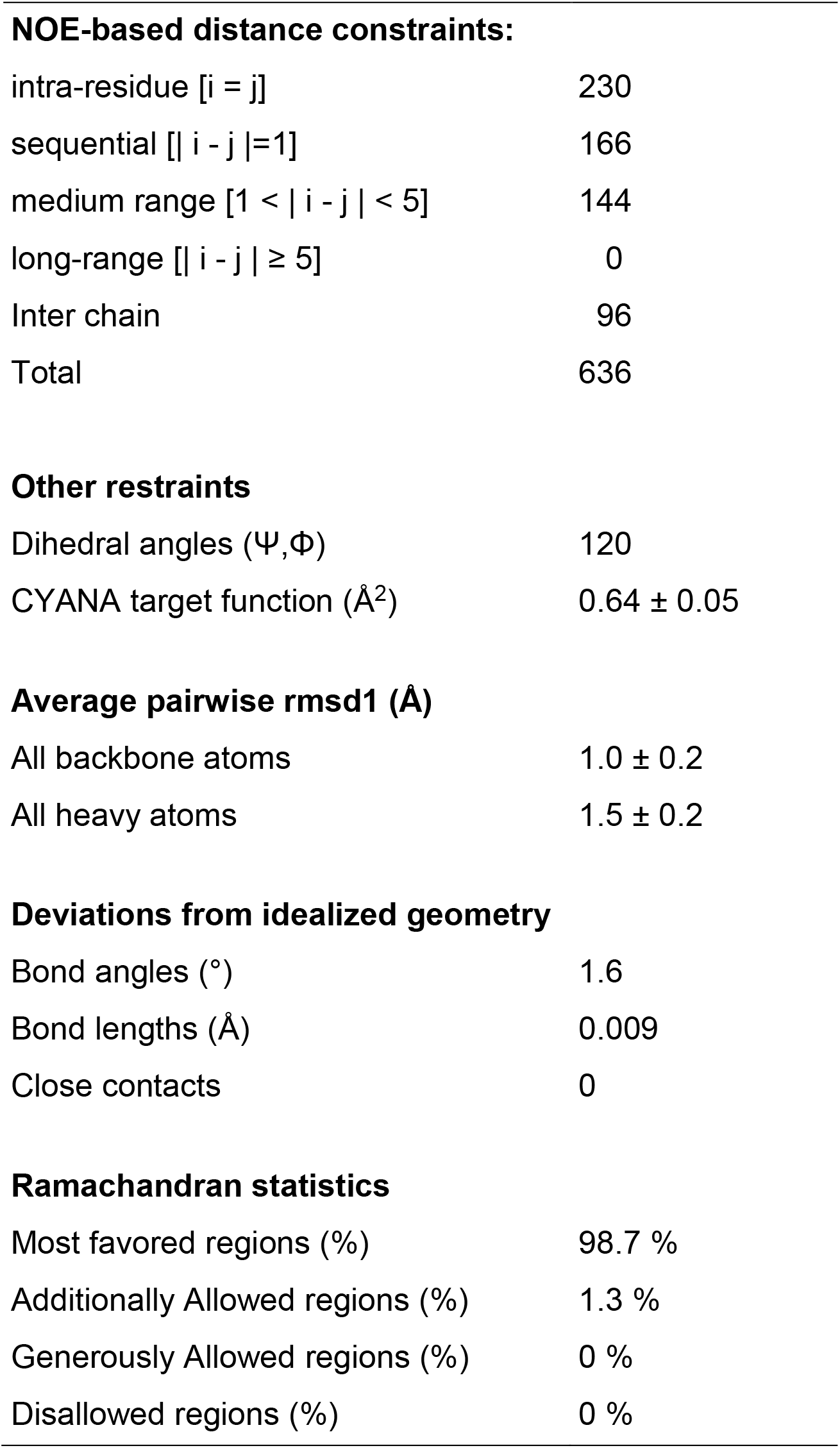
NMR and refinement statistics of human Palb2cc.

### Molecular dynamics simulation protocol

The NMR structure of Palb2cc was used to carry out unbiased molecular dynamics simulations. All simulations in this study were performed in Gromacs version 5.1.2^16^ using amber03w.ff^17^. Individual mutations on Palb2cc were created by replacing each side chain from the Dunbrack rotamer library in UCSF Chimera version 1.15^18^. All the acidic and basic residues were modeled in their charged states. The initial structures were solvated using the TIP4P2005 water model in an appropriate box. The net charge in the system was neutralized by adding an appropriate number of Na+ and Cl-ions. The neutralized system was subjected to energy minimization using the steepest descent minimization in a maximum of 50000 steps until the maximum force on each atom was less than 1000 kJ/mol/nm. The energy-minimized systems were further equilibrated under constant temperature (NVT) for 100 ps and constant temperature and pressure (NPT) for 100 ps. The temperature was maintained at 300K using a V-rescale Berendsen thermostat with a time constant of 0.1 ps. The pressure was held at 1 bar by employing the Parrinello-Rahman barostat with a time constant of 2 ps. The final production runs were carried out under periodic boundary conditions for 500 ns with a 2 fs time step. The van der Waals and short-range electrostatic interactions were calculated using a 1 nm cutoff. The long-range electrostatics were computed using the Particle Mesh Ewald method. All bond lengths were constrained using the LINCS algorithm. RMSF, RG, and RMSD were calculated using scripts available in Gromacs. Percentage ɑ-helicity was calculated using the DSSP tool in Gromacs. The trajectories were converted to amber format using Pamed. Native contacts for each mutant were computed using the CPPTRAJ version 4.14.0^19^(AmberTools V19.11). The data was plotted using in-house scripts in Python.

The phosphoserine Palb2 variant were modelled in UCSF chimera version 1.15 by replacing S10 and S29 with phosphoserines. An inhouse modified amber03w.ff forcefield, which included parameters for phosphoserine, was used to carry out the Palb2 homodimer simulations. The Palb2-BRCA1 structure was modeled by the Swiss model using PDB: 7k3s as a template. The model was then simulated for 100 ns, and the final structure was used for further simulations. A similar protocol as the Palb2 homodimer was followed to simulate the Palb2-BRCA1 heterodimer and the variants of Palb2 in complex with BRCA1. Data analysis was performed similarly.

## Results

### Palb2 coiled-coil domain forms a homodimer

Palb2cc (residue 6-41) was expressed in *E. coli* and purified to evaluate the secondary structure. The secondary structure calculated by Circular Dichroism (CD) spectroscopy revealed a highly helical (88%) segment (Figure 1C), which is consistent with the predicted coiled-coil domain. Palb2 was studied by analytical size exclusion chromatography to study the oligomerization properties. The protein eluted as a single peak corresponding to the dimer size of 8.9 kDa (Figure 1D).

### Structure of the Palb2 coiled-coil domain

The structure of the human Palb2 coiled-coil homodimer was determined using a uniformly labeled ^13^C, ^15^N sample. 0.5 mM sample was dissolved in 50 mM Na_2_HPO_4_, 150 mM NaCl, 5 mM DTT, and pH 6.5 buffer and supplemented with 10% D_2_O. All NMR experiments were collected at 25℃ using Bruker ASCEND III 600 and 800 MHz NMR spectrometers. Conventional triple resonance experiments were performed to determine the backbone and sidechain assignments (Figure 2A). Dihedral angles were calculated using the backbone assignments. The distance constraints were generated from ^15^N-NOESY-HSQC and ^13^C-NOESY-HSQC experiments with a mixing time of 100 ms. ^13^C-filtered NOESY experiments were used to determine the Palb2cc dimer interface. The distance constraints from all NOE experiments and the backbone dihedral angles were used to calculate the solution Palb2cc homodimer structure. The NMR and refinement statistics are provided in Table 1. The structure was further equilibrated in a water box before analysis.

**Figure 2.**
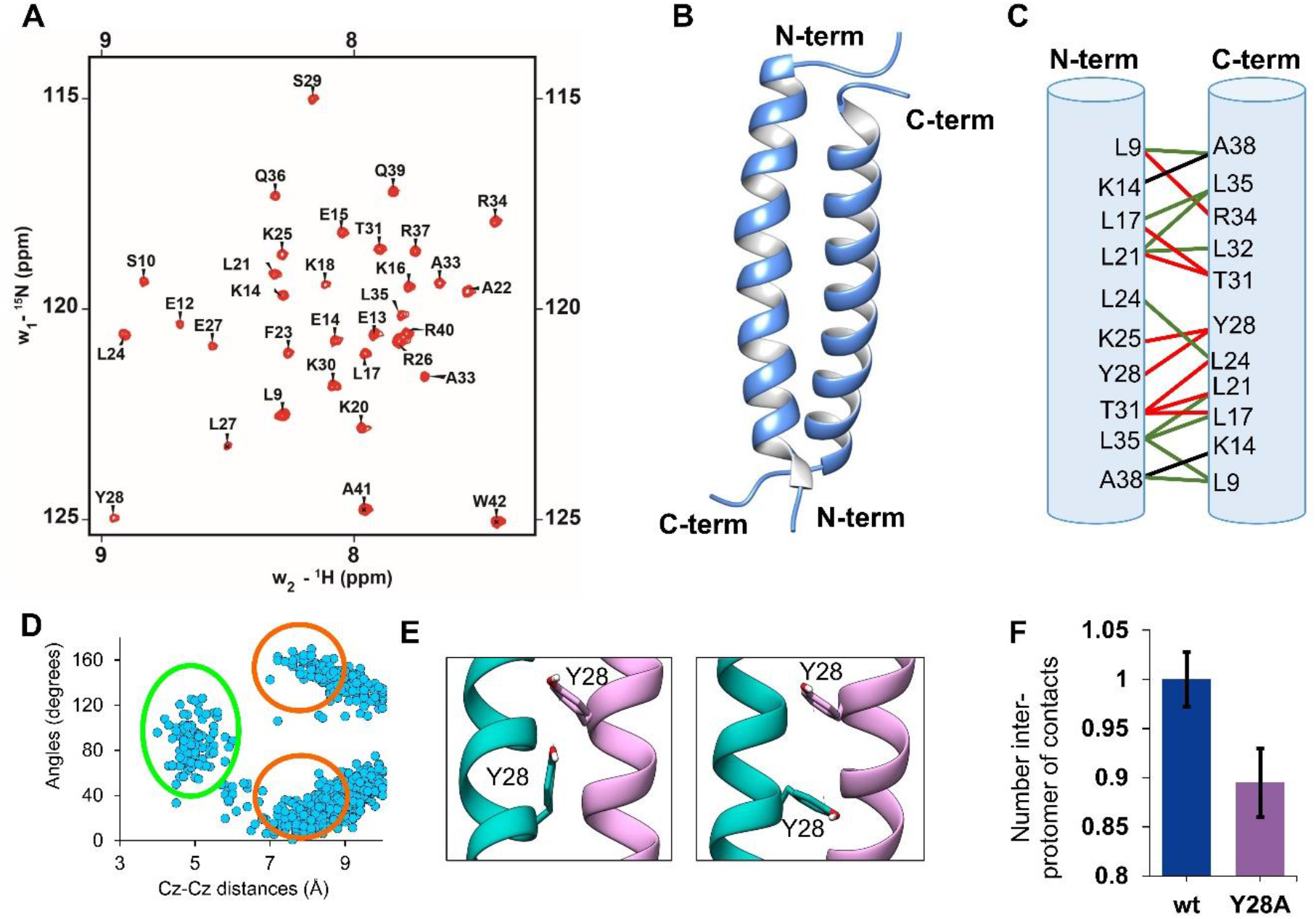
(A) The ^1^H-^15^N HSQC spectrum of Palb2cc domain (residues:6-41). The assigned backbone N-H amide resonances are labeled. The amide peaks of the first two N-terminal residues (residue numbers 6 and 7) are broadened due to exchange, signifying a dynamic region. Residue number 8 is a proline whose amide peak is not visible in the HSQC. (B) The solution structure for the Palb2cc domain suggests an antiparallel orientation between the two helices. (C) The contacts between protomers in the Palb2cc dimer are highlighted. Green, black, and red lines connect the interacting residues. The green lines denote hydrophobic contacts between leucines and alanines. Black lines show the hydrogen bonds and the red line shows the Van der Waals interactions. (D) The angle between the aromatic rings of Y28 from each protomer is plotted against the distance between C_z_-C_z_ atom pairs. The cluster of structures with edge-to-face orientation are marked in green and parallel orientation are marked in red. (E) Snapshots from the MD simulations shows the edge-to-face and paralle orientation of the aromatic rings of Y28. (F) The number of inter-protomer contacts (normalized) in the wt and Y28A variants are plotted.

The structure of Palb2cc suggests that the domain forms an antiparallel coiled-coil dimer, where the helix in each monomer stretches from residue 11 to 39 (Figure 2B). The buried surface area is 6020 Å^2^, suggesting an extensive and stable interface. A robust hydrophobic interaction network holds together the coiled-coil dimer and resembles a leucine zipper structure. The side chains of hydrophobic residues (L9, L17, L21, L24, Y28, T31, and L35) interact across the dimer interface (Figure 3C). At the center of the helices, L24 and Y28 form multiple interactions. Towards the terminal ends, R34 interacts with both K16 and K20. In addition, T31 forms contacts with L17 and L21. L35 packs against L9, L17 and L21. The human Palb2cc domain has 87% sequence identity to the mouse Palb2cc domain (Figure S1A). The structure of the human Palb2cc domain is similar to the antiparallel orientation of the mouse Palb2cc domain^20^(PDB:6e4h) (Figure S1B).

**Figure 3.**
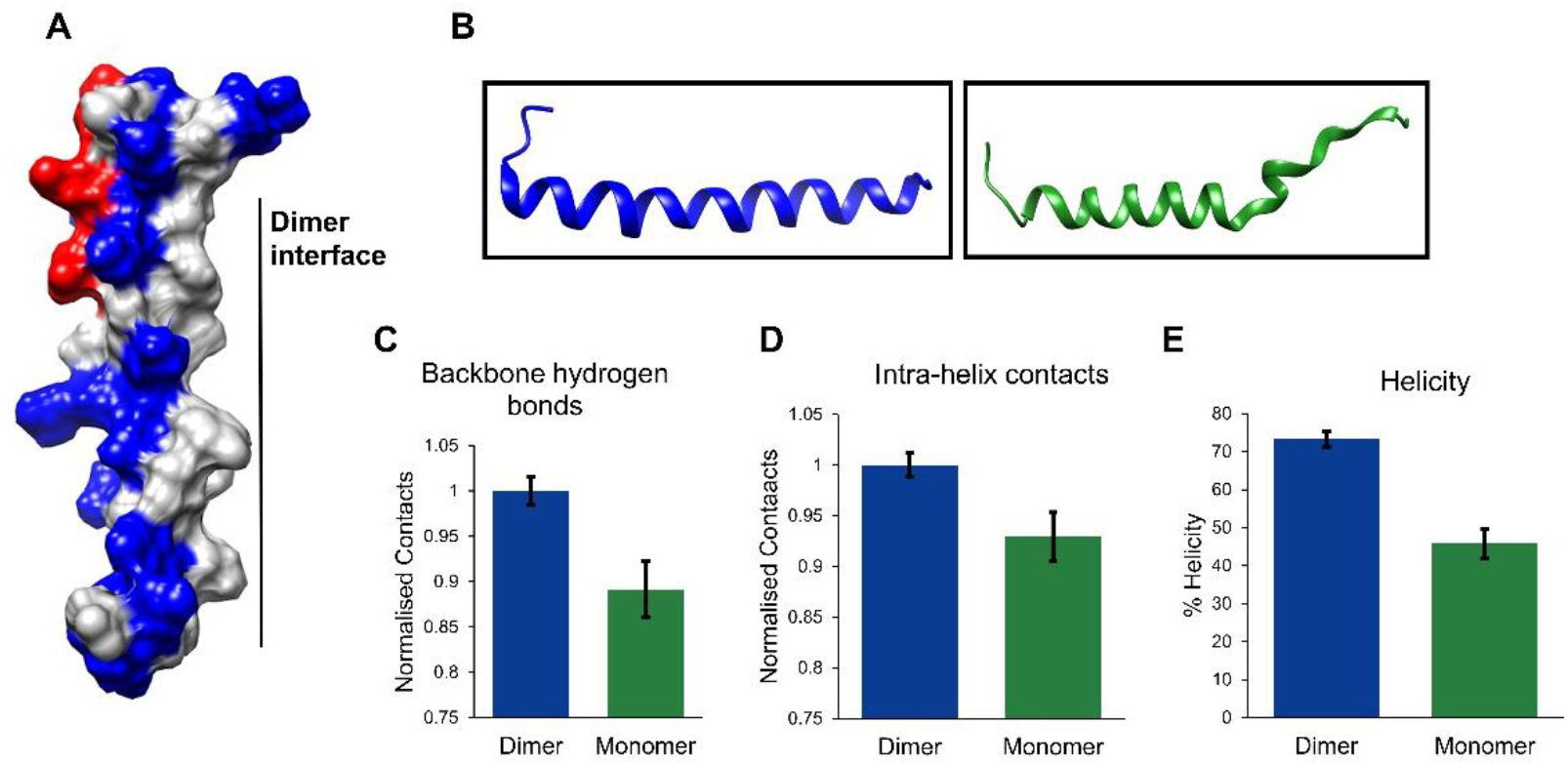
The surface charge distribution and stability of the Palb2cc domain. (A) Charge distribution on the surface representation of Palb2cc monomer (Red – Negatively charged, Blue –Positively charged, Grey-uncharged) (B) Snapshots of a single helix from Palb2cc dimer simulation (blue) and monomer simulation (green). The snapshot(C) The number of backbone hydrogen bonds in the Palb2cc dimer and monomer. (D) Intra-protomer contacts are plotted for the dimer and monomer. The contacts are measured between all non-hydrogen atoms with a distance cutoff of 7 Å. (E) Percentage helicity.

### Contacts at the dimer interface stabilize the Palb2cc protomers

Aromatic ring geometry is known to play an essential role in determining the fold and topology of the protein. Studies on structural analysis of aromatic pi-pi interactions in several proteins have shown the preference for either edge-to-face or parallel displaced stacking interaction^21^. We noticed an aromatic pi-pi interaction between the tyrosine residues (Y28) at the center of the homodimer interface. To analyze the tyrosine interactions, the Palb2cc was simulated by molecular dynamics simulations for 1.5 μs (0.5 μs x 3). The simulations of the Palb2cc domain show that the hydroxy benzene ring of Y28 from both units forming parallel orientation, and transiently forming edge-to-face orientation (Figure 2D, 2E). These interactions may strengthen the interaction network to stabilize the dimer. It has been reported that the Y28A mutant compromises PALB2 function in HR repair and produces hypersensitivity to mitomycin C treatment^22^. We then simulated a variant of Y28A-Palb2cc, which showed a reduction in the inter-protomer contacts (Figure 2F). There was a drop in the intra-helix contacts (Figure S2), suggesting that Y28 pi-pi interaction could significantly stabilize the protomer.

To study the stability of Palb2cc dimers and monomers, a total of 1.5 μs (0.5 μs x 3) MD simulations were run for Palb2cc dimers and monomers using the Amberff03 force field. The helicity of the Palb2cc was maintained throughout the simulation in the dimer. The leucine-rich dimer interface, along with Y28 and T31, maintains the structural integrity of the dimer. In the central region of the helix, L24 from both the monomer units pack against each other. Flanking the central region, the leucine residues L17 and L21 interact with L35 and L32. They create a hydrophobic pocket that is excluded from the solvent.

One side of the Palb2cc domain helix is leucine-rich and hydrophobic, essential for dimerization (Figure 3A). The other side of the helix is hydrophilic and has multiple charged amino acids. Due to the numerous lysine and arginine residues on the hydrophilic side, the coiled-coil domain carries a net positive charge. There is an uneven charge distribution along the N-C axis of the helix. The C-terminus is positively charged, as shown in Figure 3A, while the N-terminus has an equal distribution of positively and negatively charged residues. The Cα Root Mean Square Fluctuations (RMSF) describe the residue-wise fluctuations observed during the simulations. The C∝ RMSF values were higher in the individual monomer than in the dimer (Figure S3A). When an individual monomer of the Palb2cc was simulated, the monomer unfolded within 0.1 μs (Figure 3B, S3B, and S3C). This contrasts sharply with the dimer, which was stable for the entire time. The backbone hydrogen bonds, the total intra-protomer contacts, and the helicity were also reduced in the monomer than the dimer (Figure 3C-3E), suggesting that the packing of hydrophobic contacts at the dimer interface is essential to maintaining the structural integrity of each unit. Each monomer unit is prone to disorder without the interaction energy from the interfacial contacts.

### L35P mutation destabilizes the Palb2cc helix

L35P is a VUS in *Palb2* that may be associated with increased risk of breast cancer. Functional assays using mouse embryonic stem cells from *Palb2* knockout mice suggested that L35P mutation reduces HR function by more than 90% ^23^. It is unclear how the mutation destabilizes Palb2 dimerization and/or inhibits its interaction with BRCA1. To understand the structural implications of the L35P mutation on the Palb2cc domain, the secondary structure was determined by CD spectroscopy. The helicity in L35P-Palb2cc (25% helical) was reduced compared to wt (88% helical) (Figure 4A). When the molecular weights were determined by analytical size exclusion chromatography, the average molecular mass was reduced from 8.9 kDa for wt-Palb2cc to 6.8 kDa for L35P-Palb2cc (Figure S4A). The reduced molecular weight implies that the L35P substitution reduces the affinity of Palb2cc dimer, and the protein population is now a mixture of dimers and monomers.

**Figure 4.**
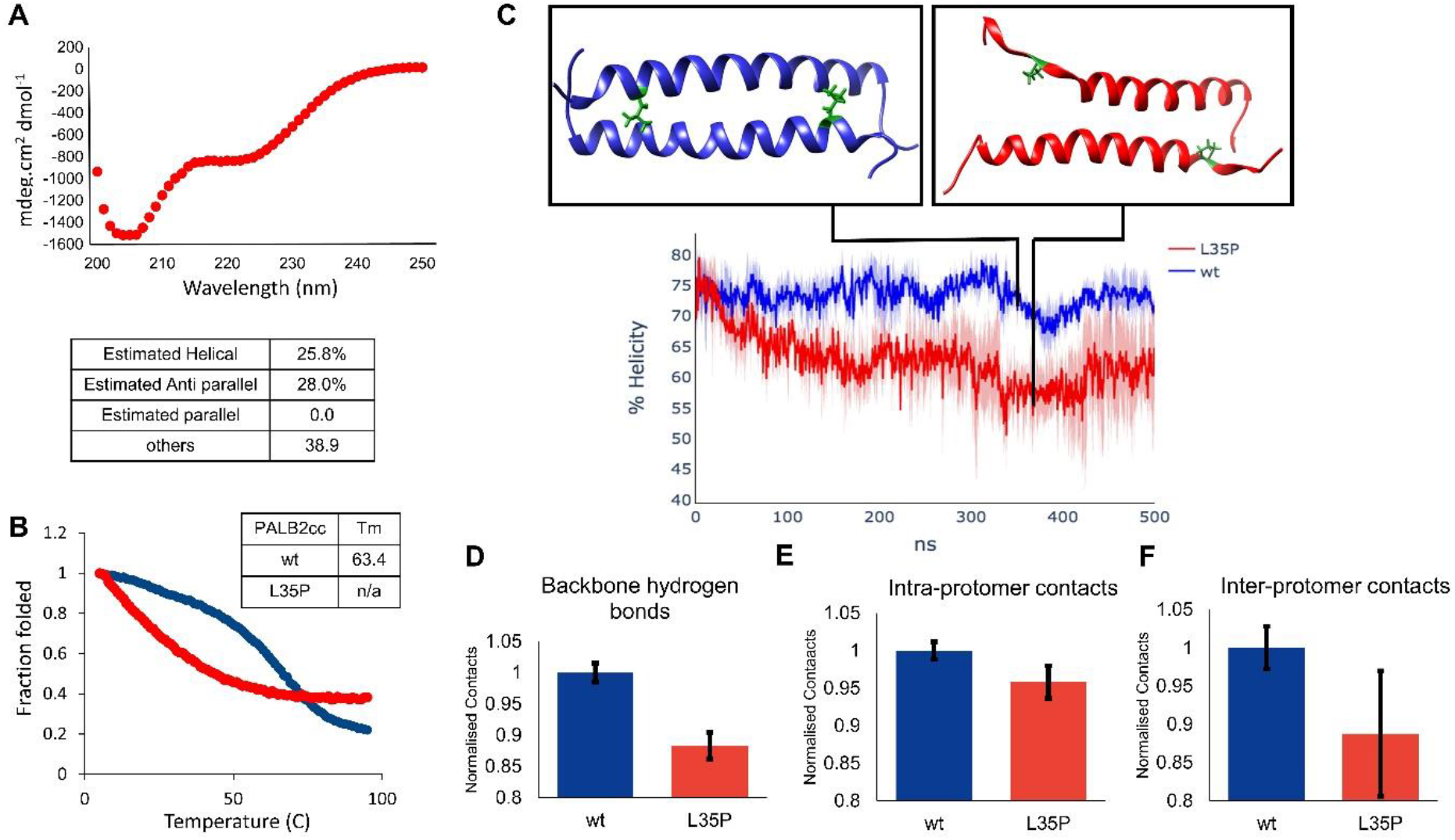
(A) The CD spectrum of L35P mutation in the Palb2cc domain. The % helicity is provided below. (B) A thermal melt of wt-Palb2cc and L35P-Palb2cc. (C) Snapshots of the wt-Palb2cc dimer (blue) and L35P-Palb2cc (green) from MD simulations. (D) The number of backbone hydrogen bonds in wt-Palb2cc and L35P-Palb2cc. (E) The number of intra-protomer contacts between all non-hydrogen atoms within the distance of 7 Å is plotted for wt-Palb2cc and L35P-Palb2cc. (F) The number of intra-protomer contacts between all non-hydrogen atoms is plotted for wt-Palb2cc and L35P-Palb2cc.

The stability of the wt-Palb2cc and L35P-Palb2cc was probed by thermal denaturation melt. The thermal stability melting point (T_m_) of wt-Palb2cc was 63.4 ºC. L35P-Palb2cc started unfolding at lower temperatures, suggesting a higher propensity to form disordered structures (Figure 4B). The melting profile of L35P-Palb2cc had a non-cooperative signature, suggesting partially unfolded intermediate states that are in equilibrium with the native and unfolded states. A 2D ^1^H-^15^N HSQC for the L35P-Palb2cc mutant was recorded and overlaid with the wt-Palb2cc domain. The backbone amide proton (^1^H) resonance chemical shift for L35P-Palb2cc spanned a narrower range than the wt-Palb2cc domain, indicating a disordered protein (Figure S4B).

To further probe the consequence of the L35P mutation on the Palb2cc structure, 1.5 µs MD simulations were performed for the L35P mutation. The C-terminus of the Palb2cc unfolded during the simulation (Figure 4C). Residues around the site of the L35P mutation showed a more significant change in Cα RMSF than wt-Palb2cc, indicating the destabilizing effect of the mutation on the helix (Figure S4C). The percentage helicity (ɑ-helix) of wt and L35P mutant were calculated using the DSSP tool in Gromacs. The helicity of wt-Palb2 was 74% during simulation, but the L35P mutation decreased the helicity to 62% within 200 ns (Figure 4C). The loss of backbone hydrogen bonds and intra-protomer bonds suggest the partial loss of secondary structure (Figures 4D and 4E), commensurate with the CD spectroscopy data. Concomitantly, the interprotomer contacts were reduced in the mutant (Figure 4F), reflecting the reduced affinity between dimers observed in the ASEC experiment.

### Relevance of genetic variants to the structure of Palb2cc homodimer

The variants L24S, R37H, and Y28C in Palb2cc reduce HR by >60%. In contrast, K18R and T31I do not significantly affect HR ^8,9^. We simulated all five mutants independently to understand the effect of each mutation on the Palb2cc homodimer. L24 of the two monomers pack against each other at the Palb2cc dimer interface with the Cδ-Cδ distance between the leucine sidechain being close to 4 Å. Substitution of L24 to A24 disrupts the dimer interface, and the mutant remains a monomer ^20^. Similarly, L24S reduced the inter-protomer contacts in the MD simulations, destabilizing the dimer (Figure S5 and S6). In addition, intra-protomer backbone hydrogen bonds were reduced significantly. The contacts of L24 were reduced by 40% when substituted with S24.

R37 did not form substantial intra-or inter-protomer interactions. Substitution of R with H does not significantly affect the charge distribution in the region. In the MD simulations, R37H increases in intra-protomer contacts and maintains similar inter-protomer contacts (Figure S5). Hence, R37H may not destabilize the Palb2cc dimer. Y28 is at the center of the Palb2cc helix. Since the hydrophobicity of Cysteines is high, the hydrophobicity at the center is maintained in Y28C. However, cysteine has a smaller sidechain than tyrosine, and multiple contacts of Y28 are disrupted in the mutant (Figure 5B). Moreover, substituting the tyrosine will disrupt the aromatic pi-pi interaction to destabilizing the homodimer. Overall, the Y28C substantially loses inter-protomer contacts, which might destabilize the Palb2cc helix, making it non-functional (Figure S6). The substitution K18R decreases inter-protomer contacts severely, thus reducing the Palb2 dimer stability (Figure S6). The sidechain of T31 interacts with the hydrophobic pocket formed by L17, L21, and L24 from the other monomer and L35 from the same chain. Even though this is an interface mutation, the hydrophobicity and sidechain geometry are maintained. Consequently, I31 and inter-protomer contacts are conserved (Figures 5 and S6). The isoleucine replaces the threonine without disrupting the interface, keeping the Palb2cc intact.

**Figure 5.**
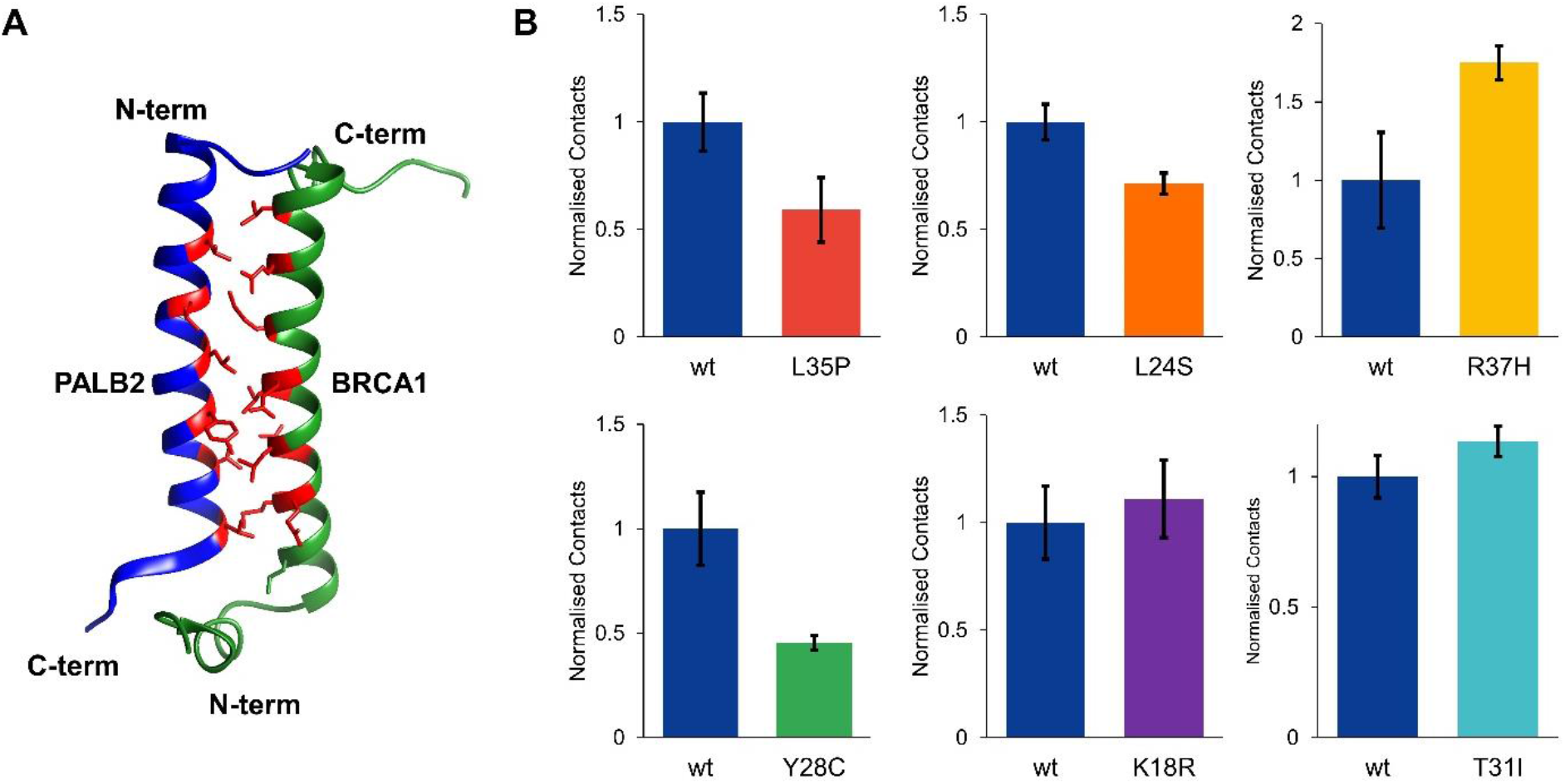
(A) Modeled with Swiss model after 100ns simulation (Blue-Palb2cc, Green-BRCA1cc, Red-dimerization interface residues). (B) Palb2cc/BRCA1 contacts at the site of Palb2 mutation. Native contacts are calculated between non-hydrogen atoms with a pairwise distance cutoff of 10 Å.

Palb2 is phosphorylated at the N-terminus during stress conditions^2^. Ionization radiation induces phosphorylation of Palb2 at serines S59, S157, and S376, which modulates its activity ^2^. The coiled-coil domain of Palb2 has two serine residues (S10 and S29) that could be potentially phosphorylated under different stress conditions. S10 is situated at the N-terminal disordered region in Palb2cc, whereas S29 is positioned at the center of the domain. We investigated whether phosphorylation at these serine residues could affect the structure of Palb2. To mimic phosphorylation, we substituted the serine residues with glutamic acid in the domain. The CD spectra and size-exclusion profile of S10E, S29E-Palb2cc, and wt-Palb2cc are similar, suggesting that the secondary structure and size of the domain are unperturbed (Figure S7A, B). We also performed MD simulations of Palb2 after replacing the serine sidechains with phosphoserines. We found that the intra- and inter-helix contacts in the phosphorylated Palb2 coiled-coil domain were similar to those in the wild-type (Figure S7), suggesting that phosphorylation may not affect the structure of the coiled-coil domain.

### Structural effects of the Palb2 variants on Palb2-BRCA1 heterodimer

We investigated the structural properties of the Palb2 coiled-coil domain in complex with the BRCA1 coiled-coil domain. To achieve this, we created a heterodimeric model using a previously reported structure of the mouse Palb2-BRCA1 (PDB: 7K3S). The model was equilibrated for 0.1 μs, and the final structure was used for further simulations for 1.5 μs (0.5 μs x 3). Throughout the entire simulation time, the Palb2-BRCA1 complex remained stable, with both protomers retaining their helicity. The Palb2-BRCA1 heterodimer retains a similar dimer interface as the Palb2 homodimer. The angle between the two helices is 24 degrees. The BRCA1cc domain has a hydrophobic interface that, like the Palb2cc domain, consists of multiple leucine-rich residues. Additionally, the BRCA1cc domain has four positively charged residues and seven negatively charged residues, creating an overall net negative charge. In particular, the C-terminal end of the helix is highly negatively charged. The positively charged N-terminal region of the Palb2cc domain allows for good charge complementarity in the heterodimer interface (Figure S8).

The hydrophobic residues of Palb2 pack against the hydrophobic face on the BRCA1 coiled-coil domain. The residues L9, L17, L21, L24, Y28, T31, L32 and L35 of Palb2cc domain engage in multiple contacts with L1392, M1400, L1404, L1407, M1411, L1414 and L1418 residues of BRCA1. Multiple leucine residues on the BRCA1 interface allow for a similar intricate hydrophobic network as observed in the Palb2 homodimer (Figure S9A). Y28 aromatic ring of Palb2 is close to M1411 of BRCA1, forming strong Met-Aromatic interactions between the two protomers ^24^. On the other side of the Y28, L1404 from the BRCA1 contacts the aromatic ring, forming CH-Pi interactions (Figure S9B). In addition to the leucine zipper between the protomers, both these interactions could enhance the stability and strengthen the heterodimeric complex.

To gain further structural insights into the effects of Palb2 variants on the Palb2-BRCA1 interactions, we performed similar simulations as described previously. L35P reduces helicity and interprotomer contacts in the Palb2 homodimer. About 50% loss in interfacial contacts is observed in the L35P Palb2-BRCA1 heterodimer (Figure 5B). A moderate reduction in native contacts was observed at the mutation site for the L24S and Y28C Palb2 variants. L24 is at the center of the helix, forming multiple interactions with leucine residues of BRCA1. A swap with a polar residue like serine disrupts the native contacts with the BRCA1 up to 30%. Y28C reduces the size of the amino acid, causing loss of Met-Aromatic and CH-Pi interaction of the Y28 aromatic ring (Figure S9C-D), overall disrupting the native contacts network below 50%, as seen in Figure 5B.

T31I, R37H, and K18R mutants showed a slight increase in the number of contacts at the mutation site. This is either due to increased sidechain size (K18R) or a change in the geometry or polarity (R37H, T31I). The K18R variant establishes a stronger charged interaction with D1419 from BRCA1. A 20% increase in the occupancy of this salt bridge is observed (Figure S10). R37 has an electrostatic interaction with D1381, S1387, and D1390 in the Palb2/BRCA1 heterodimers. This allows for partial stabilization of the unstructured N-terminus of BRCA1. H37 retains the contacts with D1381 and S1387 but has reduced interaction with the D1390. However, there is an overall gain in the total interactions observed at R37H. Overall, the L35P, L24S, and Y28C mutations substantially reduce Palb2/BRCA1 interaction, while R37H, T31I, and K18R have a higher number of contacts in the Palb2/BRCA1 complex.

## Discussion

The tumor suppressor Palb2 functions with BRCA1 and BRCA2 proteins to maintain genomic integrity by homologous recombination. Palb2 forms a homodimer by the coiled-coil domain in the inactive OFF state. Palb2 dissociates from the homodimer in the ON state during HR and interacts with BRCA1 to form a heterodimer (ON state). The switch from the OFF to ON state is a critical event in the HR response. Carriers of specific genetic variants of Palb2 have up to an eight-fold higher risk of breast cancer^25^. Two hotspots of missense variants have been identified, the N-terminal coiled-coil domain and the C-terminal WD40 domain. To understand the effect of coiled-coil domain mutations on the Palb2 structure-function relationship, we solved the human Palb2 coiled-coil domain structure. The solution NMR structure of human Palb2cc suggests that it forms an antiparallel homodimer. A network of hydrophobic interactions formed by residues L9, L17, L21, L24, Y28, L35, and A38 drives the dimerization of the Palb2cc domain. These residues in the i+3 or i+4 position create a hydrophobic patch complemented by the other protomer. The coiled-coil domain is positively charged with an uneven charge distribution from N-to C-terminal, which induces the antiparallel structure. The Palb2 structure resembles the antiparallel coiled coil formed by myosin X^26^. MD simulations indicated that the monomer coiled-coil domain unfolds rapidly compared to the dimer. The uneven charge distribution could destabilize the monomer secondary structure. Our data suggest that Palb2 homodimerization or its heterodimerization with BRCA1 is essential to the structural integrity of this domain.

Aromatic interactions are essential for protein structure and interactions ^21,27^. An analysis of aromatic interactions in protein structures shows that tyrosine is common at protein interfaces, possibly because the hydroxyl group provides stability when exposed to solvent. These interactions could be pi-pi or cation-pi interactions. Our simulations suggest that pi-pi interactions are formed between the sidechains of tyrosine 28 from the two Palb2cc protomers. The aromatic interactions between the protomers can further strengthen the hydrophobic interactions for dimerization. The reduced inter-protomer contacts and structural stability of the Y28C mutant underline the importance of aromatic interactions.

Functional assays have been performed on a few missense mutations in the Palb2cc domain. The L35P (c.104T>C) mutation was identified from a breast cancer family, a loss of function mutation that disrupts Palb2-BRCA1 interaction ^23^. Our MD simulations on the Palb2cc structure suggest the mutation completely disrupts the helical structure at the Palb2cc C-terminal end. Incorporating Proline into an α-helix is challenging due to its lack of an amide proton and the ring formed by its backbone and sidechain. Moreover, the Proline substitution disrupts multiple critical inter-protomer contacts to hamper interaction with the other protomer in the dimer, leading to a destabilized homodimer. However, the homodimer instability does not increase the heterodimer population because the mutation disrupts essential contacts with BRCA1. Overall, the L35P mutation disrupts the Palb2cc protomer structure, abolishing its self dimerisation as well as interaction with BRCA1.

The mutations Y28C, R37H, and L24S hamper HR by more than 60% and show sensitivity to the PARP inhibitor. These variants fail to pull down BRCA1, indicating a loss of binding. They also fail to localize at the site of DNA damage^8,9^. However, mechanistic insight into the mutation’s effect on the Palb2cc structure and interaction is absent. Our study shows that Y28C does not disturb intra-protomer contacts but disrupts multiple inter-protomer contacts, including the aromatic pi-pi interaction. The mutation also severely perturbs interactions with BRCA1. Hence, Y28C does not alter the monomer structure but disrupts the homo and heterodimer interactions. L24S mutation does not change the intra- and inter-protomer contacts, suggesting that the homodimer Palb2cc structure is maintained. However, it significantly disrupts BRCA1 interactions. R37H mutation shows no significant reduction in recruitment to the damaged site or interaction with BRCA1, yet causes a decrease in HR and shows sensitivity to PARP inhibitor. Possibly, R37H perturbs higher-order complex formation with BRCA proteins by allosteric mechanisms. T31I is a tolerated mutation that has minimal effects on HR. T31I does not impact Palb2cc monomeric or dimeric structure. Moreover, the functional interactions with BRCA1 are maintained in T31I, which correlates with the unaffected HR. K18R is recruited to the site of DSB more efficiently than WT. This could be due to a stronger association with BRCA1. A similar trend is observed in our simulation studies, where the Palb2 homodimer is not significantly hampered, but the Palb2/BRCA1 interactions are enhanced. The results of the structural analysis of Palb2cc variants reveal a high degree of correlation with their efficiency in homologous recombination mechanisms.

## Conclusion

Structure determination followed by molecular dynamics has suggested the effect of clinical missense mutations on the Palb2 coiled-coil domain’s fold and interactions, consistent with the mutant’s efficiency in homologous recombination. The structure and approach will help to study other variants of unknown significance in the Palb2 coiled-coil domain.

## Acknowledgments

This work was supported by the Tata Institute of Fundamental Research, Department of Atomic Energy, Government of India, under project identification no RTI 4006. The NMR data were acquired at the NCBS-TIFR NMR Facility, supported by the Department of Atomic Energy, Government of India, under project no RTI 4006. The NMR facility is also partially supported by the Department of Biotechnology, India, under project number dbt/pr12422/med/31/287/2014.

