## Supplementary information for "Structural analysis of genetic variants of the human tumor suppressor Palb2 coiled-coil domain"

```
          9                                41
Human     LSCEEKEKLKEKLAFLKREYSKTLARLQRAQRA
          ||..|||||||||||||:|||:||||||||:||
Mouse     LSYAEKEKLKEKLAFLKKEYSRTLARLQRAKRA
```

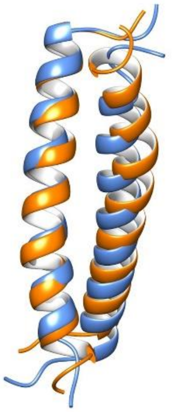

Figure S1. The comparison of primary sequence and three-dimensional structures of human (blue) and mouse (orange) Palb2cc domain.

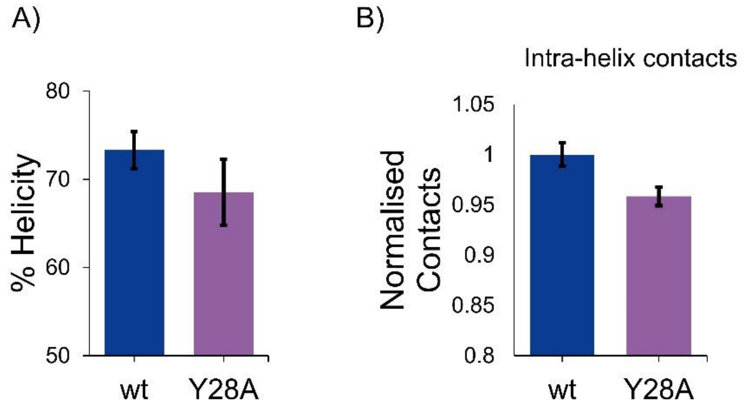


Figure S2. A) The comparison of %helicity in wt-Palb2cc and Y28A-Palb2cc. B) A comparison of intra-helix contacts in wt-Palb2cc and Y28A-Palb2cc.

A)

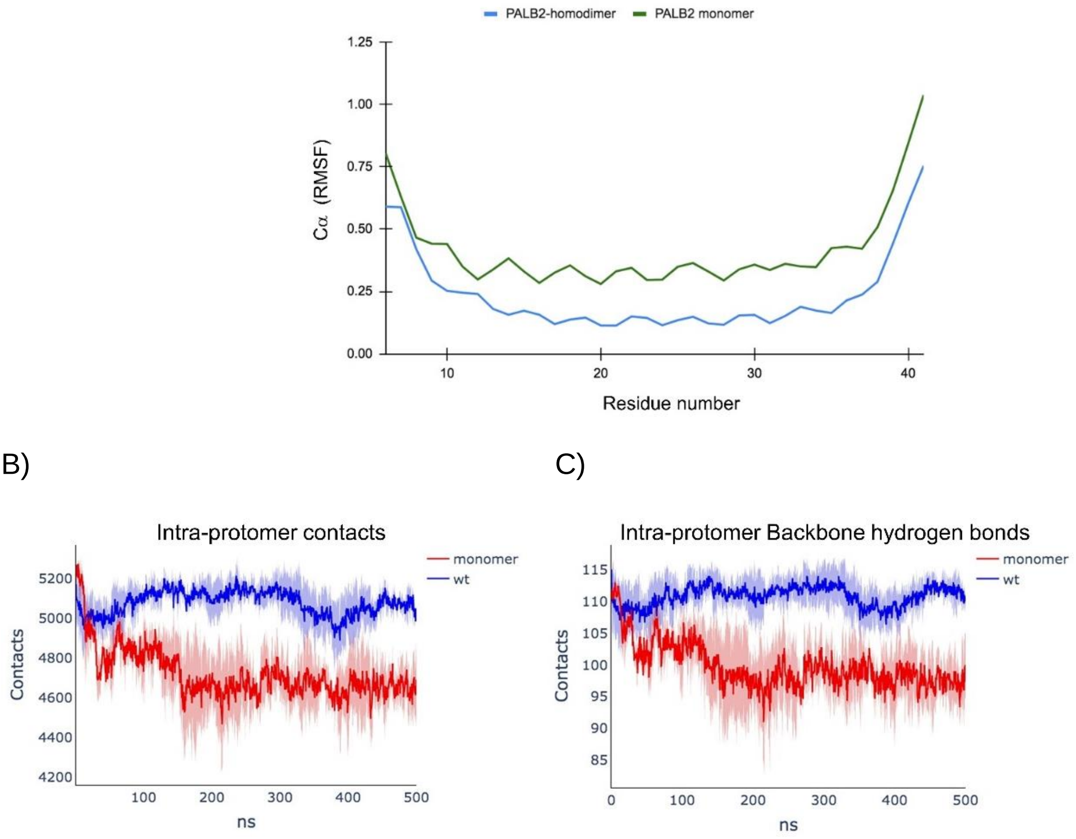


Figure S3. A) The Cα RMSF of Palb2cc homodimer (blue) and monomer (green). B) The Intra-protomer contacts and C) Intra-protomer Hbonds are calculated for the Palb2cc domain monomer. MD simulations were performed across three trajectories and the mean and std dev is plotted here.

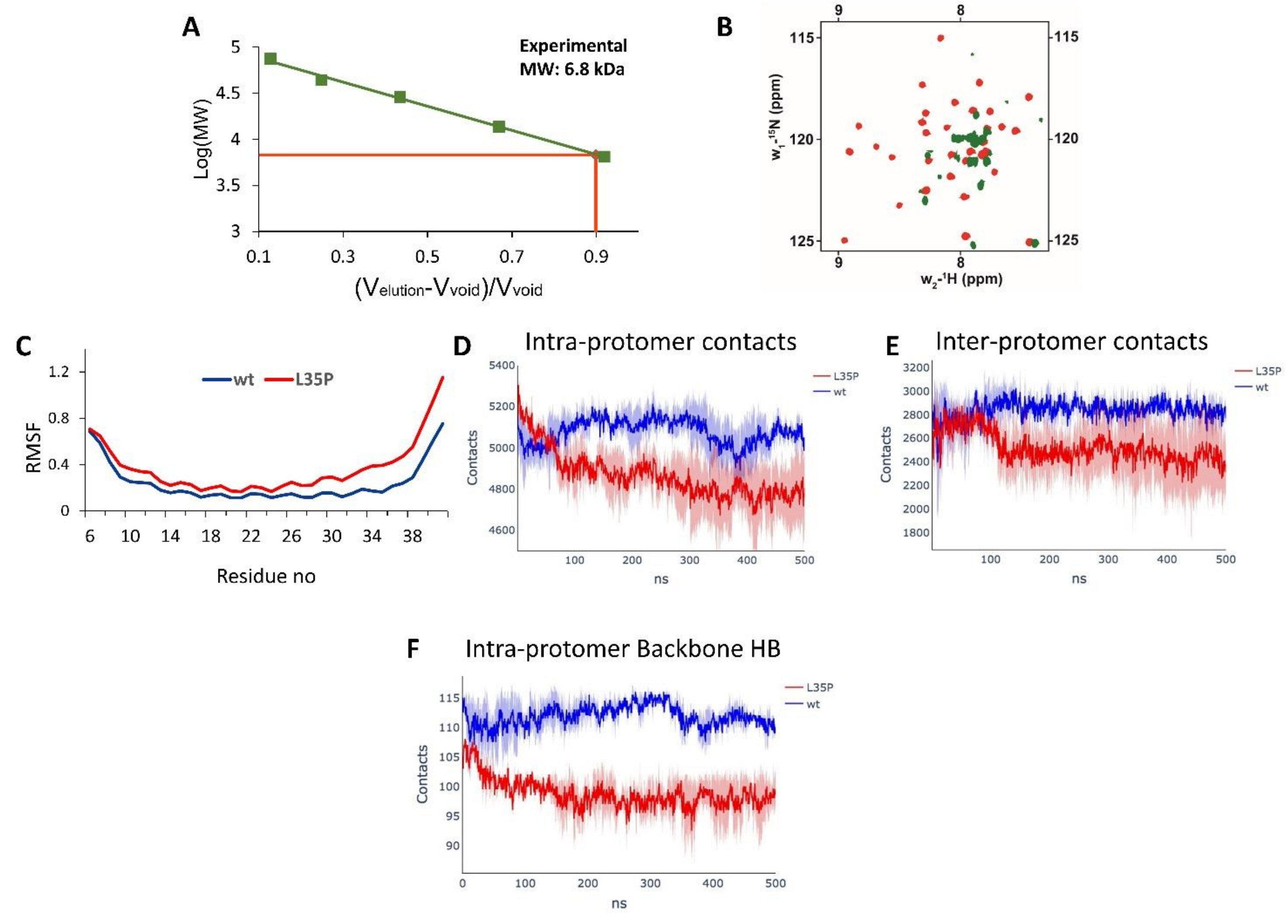


Figure S4. (A) The size of the L35P-Palb2CC domain is determined by analytical size exclusion chromatography. The plot of molecular weight standards against the elution volume is provided. The measured molecular weight is given on top of the plot. (B) The overlay of wt-Palb2cc (red) and L35P-Palb2cc (green) $^{15}$N-$^{1}$H HSQC NMR spectra. (C) The C$\alpha$ RMSF of wt-Palb2cc (blue) and L35P-Palb2cc (red). D)-F) Intra-protomer contacts, Inter-protomer contacts, and Intra-protomer Hbonds are calculated for the Palb2cc domain mutants. MD simulations were performed across three trajectories and the mean and std dev is plotted here.

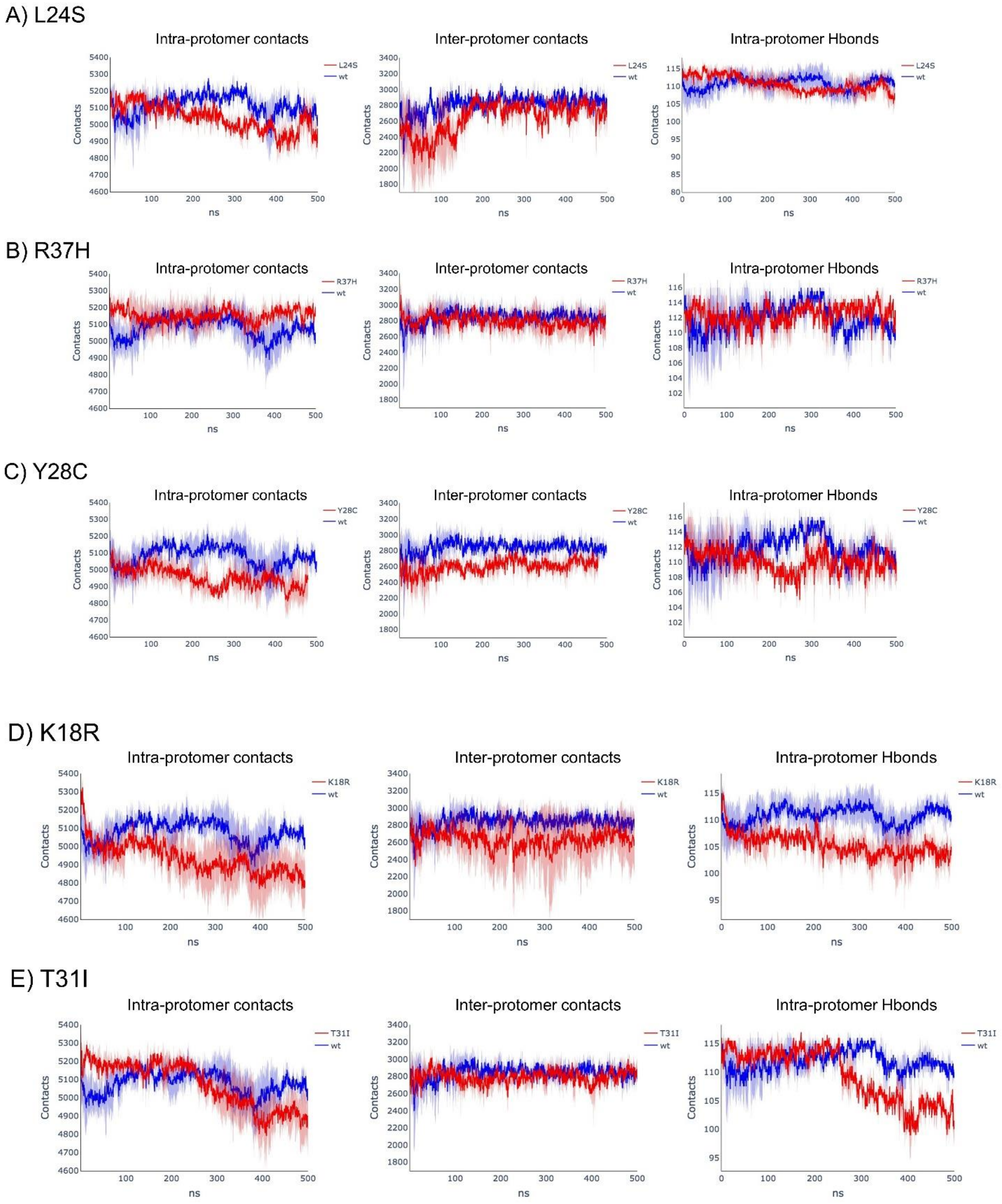


Figure S5. The Intra-protomer contacts, Inter-protomer contacts, and Intra-protomer Hbonds are calculated for the Palb2cc domain mutants a) L24S, b) R37H, c) Y28C, d) K18R, and e) T31I mutants of Palb2cc. MD simulations were performed across three trajectories and the mean and std dev is plotted here.

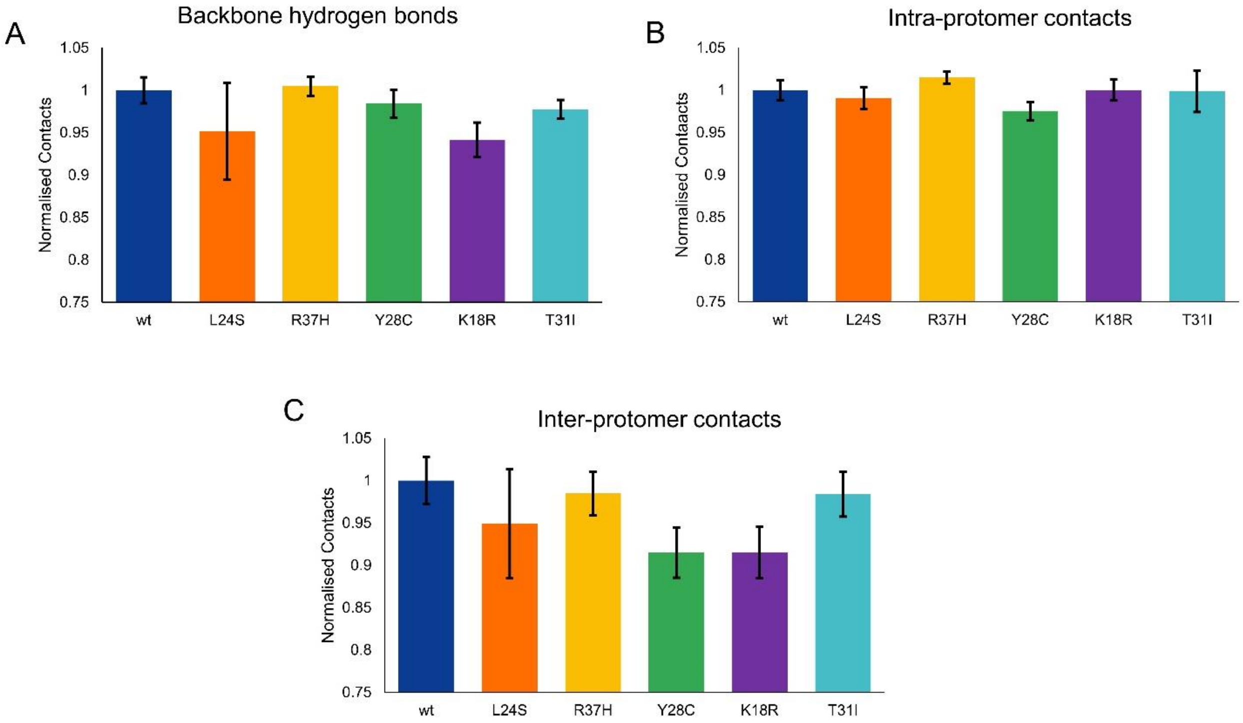


Figure S6. Comparison of L24S, R37H, Y28C, K18R, and T31I mutations in Palb2cc domain with wt (A) Backbone hydrogen bonds, (B) Non-hydrogen intra-protomer contacts (distance cutoff -7 A) (C) Non-hydrogen inter-protomer contacts (distance cutoff -10 Å).

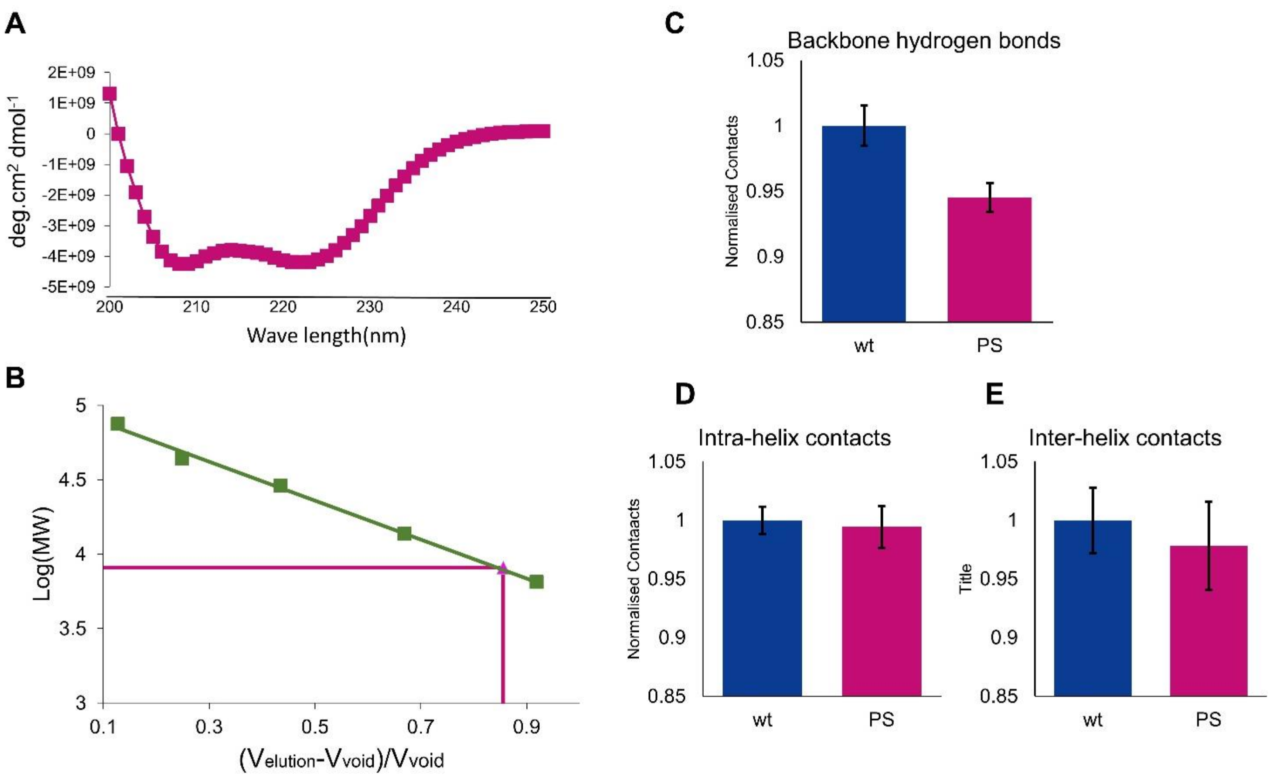


Figure S7. (A) CD spectrum of Phosphoserine mimetic mutations (S10E, S29E) mutation in Palb2 coiled-coil domain. (B) Molecular weight determination by Analytical size exclusion chromatography. A comparison of SE mutants in Palb2cc domain with wt (C) Percentage helicity (D) Number of backbone hydrogen bonds. (E) Non-hydrogen intra-protomer contacts (distance cutoff -7 A). (F) Non-hydrogen inter-protomer contacts (distance cutoff -10 A) of phosphorylated Palb2.

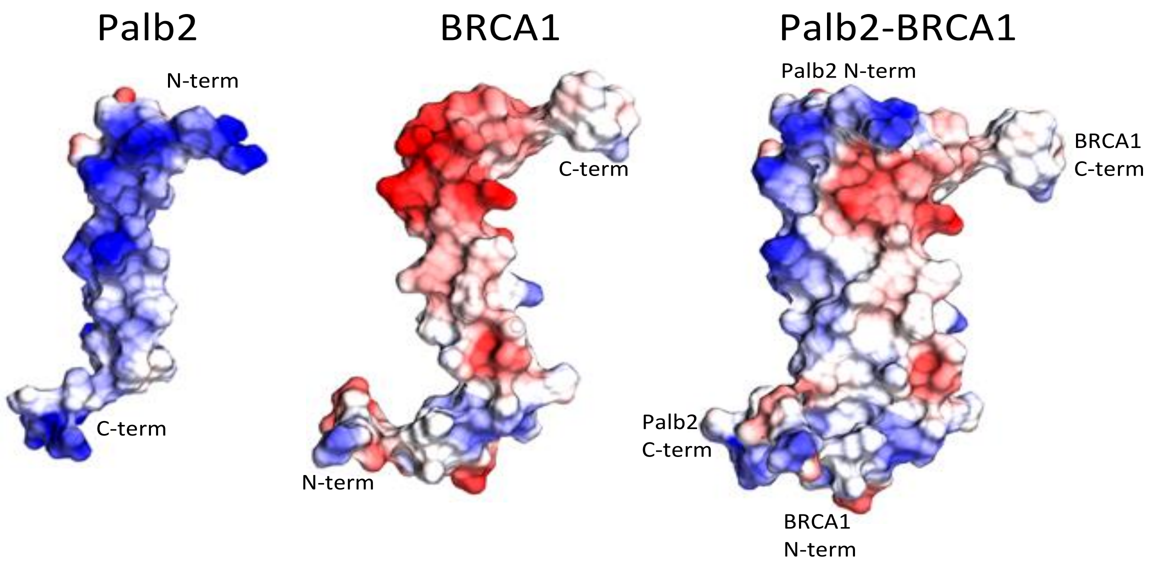


Figure S8. Columbic charge distribution on Palb2 protomer, BRCA1 protomer and Palb2-BRCA1 heterodimer complex on surface representations. (Red –Negatively charged, Blue – Positively charged, Grey- uncharged)

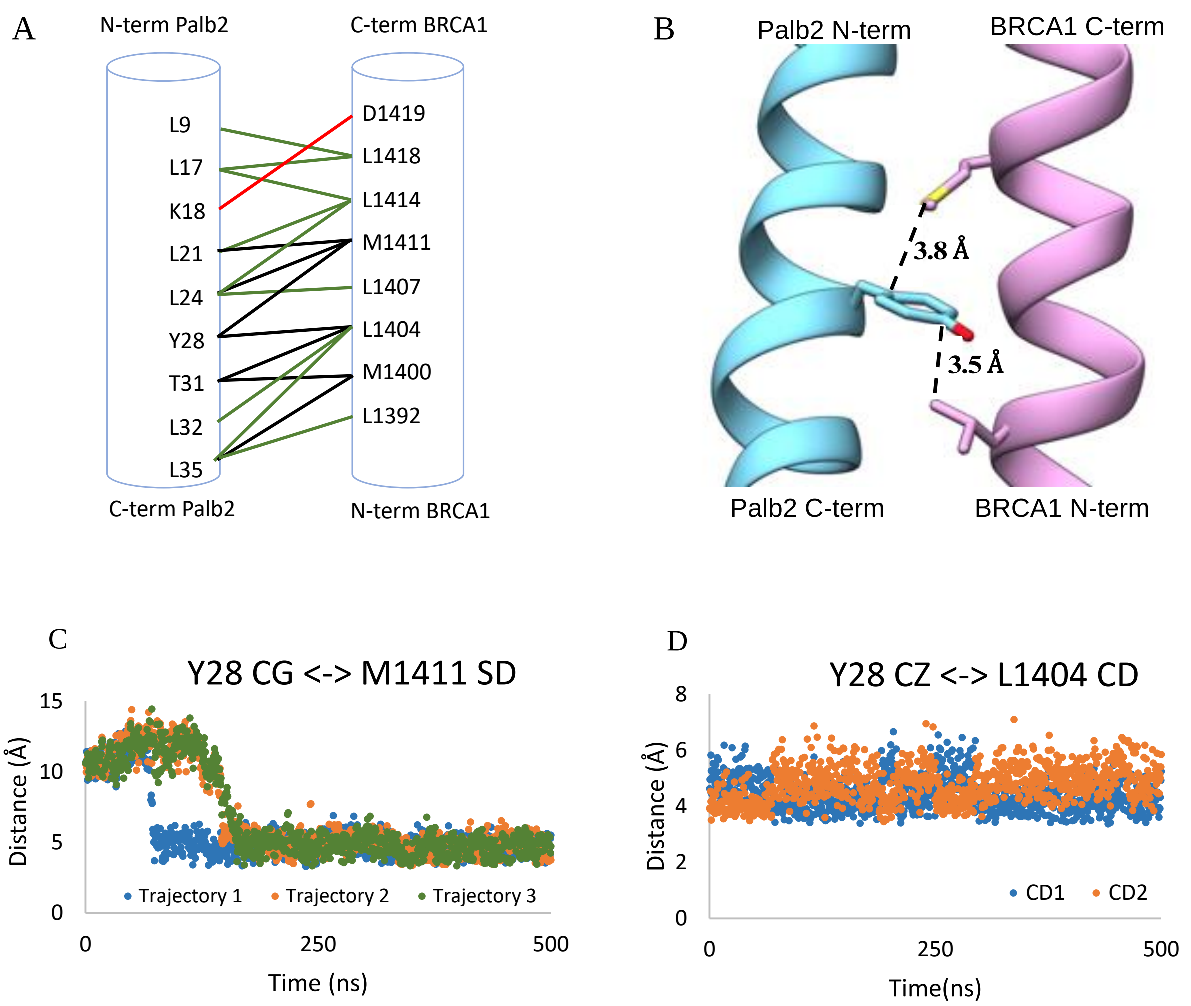


Figure S9. (A) Palb2-BRCA1 heterodimer contacts (Palb2-Blue, BRCA1- Pink). Green lines represent hydrophobic contacts, black represents van der waals interactions, red represents charged interactions. (B) Met-Pi interaction between Palb2 Y28 and BRCA1 M1411 and CH-Pi between Palb2 Y28 and BRCA1 L1404. (C) Distance between Y28 CG and M1411 SD across 3 trajectories (D) Distance between Y28 CZ and L1404 CD1 and CD2 from single trajectory.

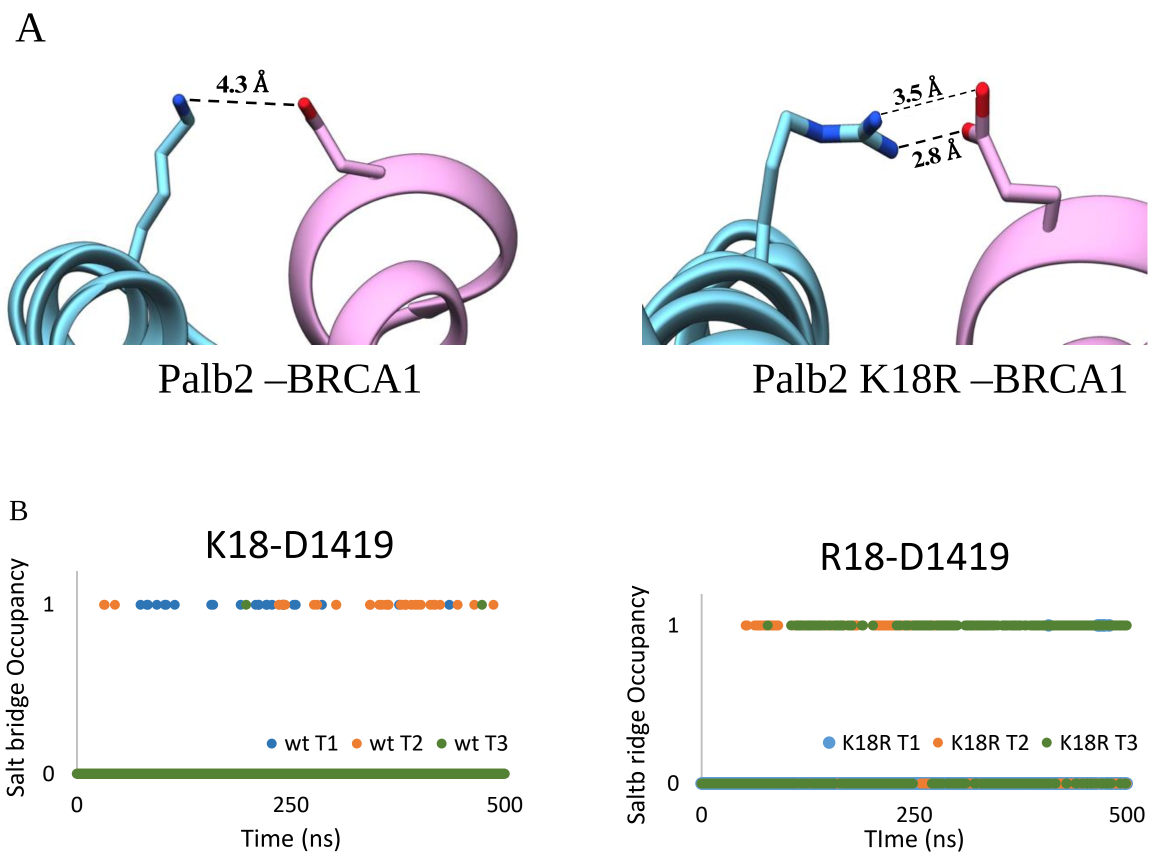


Figure S10. (A) Salt bridge between Palb2 K18/R18 and BRCA1 D1419. (B) The occupancy plot of the K18/R18 and D1419 salt bridge. N-O distances between K18/R18 and D77 sidechains were calculated and a 5Å cut-off was used to determine the occupancy of a salt bridge.
